# Adaptive generalization and efficient learning under uncertainty

**DOI:** 10.1101/2025.08.21.671603

**Authors:** Jiwon Park, Dongil Chung

## Abstract

People often use recognizable features to infer the value of novel consumables. This “generalization” strategy is known to be beneficial in stable environments, such that individuals can use previously learned rules and values in efficiently exploring new situations. However, it remains unclear whether and how individuals adjust their generalization strategy in volatile environments where previously learned information becomes obsolete. We hypothesized that individuals adaptively use generalization by continuously updating their beliefs about the credibility of the feature-based reward generalization model at each state. Our data showed that participants used generalization more when the novel environment remained consistent with the previously learned monotonic association between feature and reward, suggesting efficient utilization of prior knowledge. Against other accounts, we found that individuals incorporated an arbitration mechanism between feature-based value generalization and model-based learning based on volatility tracking. Notably, our suggested model captured differential impacts of generalization dependent on the context-volatility, such that individuals who were biased the most toward generalization showed the lowest learning errors when the value of stimuli are generalized along the recognizable feature, but showed the highest errors in a volatile environment. This work provides novel insights into the adaptive usage of generalization, orchestrating two distinctive learning mechanisms through monitoring their credibility, and highlights the potential adverse effects of overgeneralization in volatile contexts.

## Introduction

Generalization is widely recognized as an efficient decision-making strategy, enabling individuals to predict the value of novel items based on prior experiences^1,2^. It has been shown that human decision-makers can generalize the value of previously learned items to new items by comparing feature similarity across various perceptual attributes, such as color^3^, orientation^4^, and skewness^5^. However, relying solely on generalization without considering its relevance to the specific decision context can lead to poor outcomes. For instance, visually appealing mushrooms can be poisonous, and green apples may belong to different varieties rather than simply being unripe. Consequently, overgeneralization often leads to maladaptive behavior and is closely linked to various mental health disorders, including anxiety^6-11^, depression^12^, post-traumatic stress disorder^13^, and borderline personality disorder^14^. Despite the importance of calibrating generalization to fit the context given the inherent uncertainties of the real world, it remains unclear whether and how individuals adjust their generalization strategies in volatile environments, where previously learned information may no longer be reliable.

Previous studies demonstrated that individuals’ decisions are guided by various expert systems in the brain. ‘Model-free’ learning strategies enable decision-makers to rely on cached values from past experiences, while ‘model-based’ strategies involve constructing an internal model of the environment to simulate action plans and predict outcomes during decision-making^15-17^. It was suggested that the balance and arbitration between these learning (and decision-making) strategies are crucial for enabling individuals to flexibly adjust to volatile environments^16,18^. Other examples showed that such a balancing of different strategies extends to various learning contexts, consistently demonstrating that individuals primarily adopt strategies believed to be more reliable based on their experienced outcomes^19-21^. We hypothesize that individuals may apply the generalization of previously learned knowledge—a strategy typically chosen for efficient decision-making—in a similar manner, relying more on it when the environment is more stable and less so when it is volatile.

To test this hypothesis, we designed a novel probabilistic reward learning task in which participants learned the reward contingencies associated with a set of stimuli, each defined by a unique orientation. The task consisted of an initial “Anchoring session”, in which individuals were trained on three stimuli and their associated reward probabilities, followed by a “Generalization session”, where they were asked to learn the reward contingencies associated with four novel stimuli (**Fig. 1A, 1B, S1**). A total of 166 participants (male/female = 107/59, age = 23.25 ± 3.20) were randomly assigned to one of four groups, each receiving a different combination of Anchoring and Generalization sessions. For two groups of individuals, the Anchoring session was intended to implicitly convey a ‘monotonic’ positive relationship between stimulus orientation (0º, 45º, 90º) and reward probabilities (0%, 50%, 100%), whereas for the other two groups, the association was designed to be ‘non-monotonic’ (see **Methods**). There were two types of Generalization session, distinguished by the initial relationship between the reward probabilities of the stimuli and their orientation. For two groups—one that experienced a monotonic Anchoring session and one that experienced a non-monotonic Anchoring session—, the stimulus-reward association was set to be positive (‘congruent’), while for the other two groups, the association was negative (‘incongruent’). Each Generalization session comprised an initial Stable phase, during which the reward contingencies remained constant, and a Volatile phase, during which the contingencies changed unexpectedly without warning to participants (see **Methods**). We expected that the ‘monotonic-congruent (mC)’ group, compared to the other three groups (monotonic-incongruent (mIC), non-monotonic-congruent (nmC), and non-monotonic-incongruent (nmIC)), would make the greatest use of the learned experience from the Anchoring session and rely more heavily on a generalization strategy during the Generalization session, particularly during the initial Stable phase. By analyzing participants’ behavioral responses across the four groups and examining their differences, we aimed to investigate whether individuals adaptively apply generalization in different environments.

**Figure 1.**
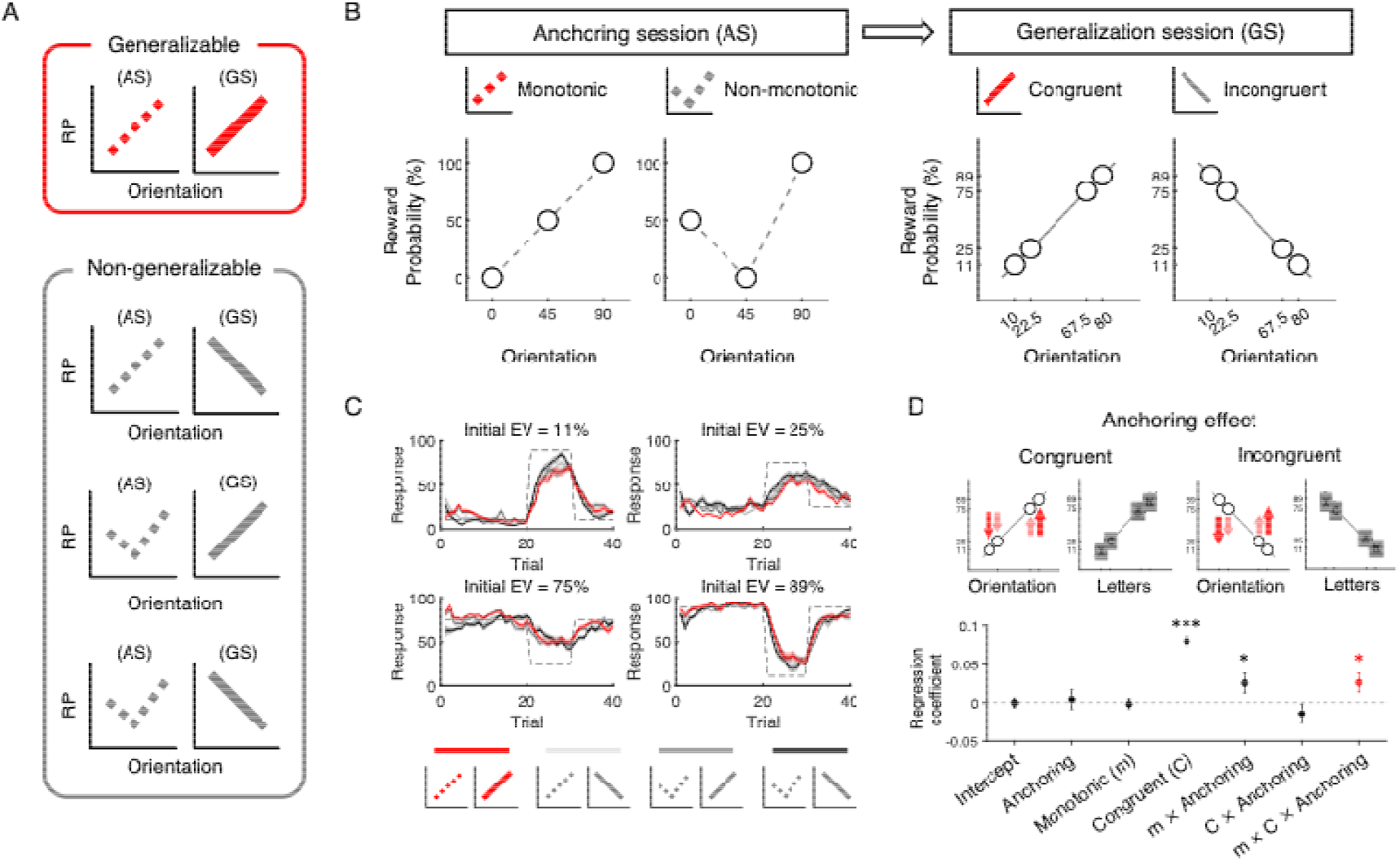
Anchoring bias and adaptive generalization in probabilistic reward learning. **(A, B)** Participants engaged in a learning task where they repeatedly observed reward outcomes to infer the reward probabilities associated with each stimulus. The task was divided into two sessions: an Anchoring session (AS) (dotted line) and a Generalization session (GS) (solid line). During the AS, participants learned either a monotonic (main diagonal) or non-monotonic (v-shape) association between stimulus orientation and its corresponding reward probabilities. In the subsequent GS, participants were introduced to four novel stimuli. The combination of a monotonic AS followed by a congruent GS (main diagonal) was designated as the generalizable context (red), whereas all other combinations with an incongruent GS (antidiagonal) were categorized as non-generalizable contexts (gray). **(C)** Reward contingencies for the stimuli were reversed twice during the task, creating a volatile learning environment. Learning curves from the GS demonstrate that participants successfully learned the reward probabilities for each unique stimulus. Dashed lines represent pre-assigned reward probabilities, solid lines show individuals’ average responses, and shaded areas indicate s.e.m. **(D)** The Anchoring effect was quantified as the tendency to overestimate the value of stimuli resembling the 90-degree orientation stimulus and underestimate the value of stimuli resembling the 0-degree orientation stimulus, compared to individuals’ value estimation for stimuli lacking generalizable features (i.e., letter stimuli instead of Gabor patches). The lower panel presents standardized regression coefficients for each regressor, with error bars representing the standard error for the standard beta estimates. \**P* < 0.05, \*\*\**P* < 0.001.

## Results

### Prior knowledge systematically biases subsequent learning in generalizable contexts

To investigate whether individuals use a generalization strategy in our reward learning task, the stimuli were designed as Gabor patches where the orientation of each stimulus could be used to infer its associated reward probability (‘main learning task’ hereafter). Given the feature similarity among stimuli, it was crucial to ensure that individuals could distinguish between them and learn the associated reward contingencies despite their resemblance. Based on individuals’ performance in both an additional orientation discrimination task (see **Supplementary text**) and the reward learning task, we confirmed that individuals across all four groups were able to distinguish all presented stimuli (**Fig. S2**) and successfully learned the assigned reward probabilities in both the Anchoring **(Fig. S3A)** and Generalization sessions **(Fig. 1C)**. During the learning task, participants were occasionally asked to re-enter the reward probability estimate they had previously reported on a given trial, serving as catch trials. The average accuracy of the these catch trials was 0.97 (SD = 0.057), across all groups.

Individuals’ baseline learning capacity, independent of the stimuli’s feature information, was also assessed using a ‘control learning task’ in which unfamiliar foreign letters (see **Methods**) were presented as stimuli instead of Gabor patches (catch trial accuracy = 0.97 ± 0.066; **Fig. S1, S3B**). By comparing individuals’ behavioral responses in the main learning task in comparison to their baseline learning capacity, we examined whether learning behaviors in the Generalization session were influenced by the learning experience in the Anchoring session. Given that the stimulus-reward relationship between the Anchoring and Generalization sessions is most consistent for the mC group, we hypothesized that generalization would be most prominent in this group as an efficient learning strategy. Specifically, if participants generalized value based on feature similarity (i.e., stimulus orientation), we expected their reward probability estimations for stimuli with lower orientations to be biased toward the pre-learned reward probability associated with the 0-degree stimulus, while those for higher orientations to be biased toward the pre-learned association with the 90-degree stimulus. To estimate the ‘Anchoring effect’—defined as the degree to which reward probabilities were systematically over- or underestimated based on stimulus orientation—, we conducted a two-step regression analysis. In the first step, residuals were computed from a regression of participants’ reward probability ratings onto the true reward probabilities in both the main and control tasks (see **Methods** for details). These residuals, reflecting orientation-based deviations, were then used as the dependent variable in a second regression. This analysis revealed that the Anchoring effect was significantly more pronounced in the mC group compared to the other three groups (m*C*Anchoring: beta = 0.026, P = 0.033; **Fig. 1D**). Note that the Anchoring effect observed in the mC group was identified after controlling for other potential main effects (C: beta = 0.080, P < 0.001; m: beta = −0.0015, P = 0.81) and for Anchoring effects shared across multiple groups (Anchoring: beta = 0.0044, P = 0.72; m*Anchoring: beta = 0.025, P = 0.039; C*Anchoring: beta = −0.014, P = 0.26). This result supports our hypothesis that prior experience influences subsequent learning, particularly to a greater extent in generalizable contexts.

### Individuals adaptively switch learning strategies based on the usefulness of prior knowledge

Given the information participants received regarding possible changes in stimuli-reward contingencies midway through the session (during the Volatile phase), an alternative learning strategy to feature-based generalization involves tracking the volatility of stimuli-associated uncertainty^22^. We hypothesized that individuals may evaluate the reliability of the generalization strategy when inferring stimulus values, and when it is deemed unreliable, switch to a volatility-tracking strategy as an alternative. To test this hypothesis, we constructed a computational model that assumes arbitration between the two learning strategies in which predictability of each strategy serve as evidence of its relative reliability compared to the alternative (**Fig. 2A**; see **Methods** for model details). A sensitivity parameter ‘*β*’ modulated how strongly the reliability difference influences arbitration. We included two additional parameters in the model to capture the baseline level of generalization. As part of the generalization strategy, the generalization width ‘*σ*’ reflected the extent of generalization—smaller *σ* indicated that only the most similar stimuli’s values were influenced by the directly experienced stimulus-reward association (see **Methods**). The parameter ‘*biasG*’ captured the bias toward using the generalization strategy over the alternative, volatility-tracking strategy (see **Methods**). These model parameters were designed to capture individual variability in generalization, which was expected to vary both within individuals over the course of learning and across different learning contexts defined by the reward contingencies assigned to each group.

**Figure 2.**
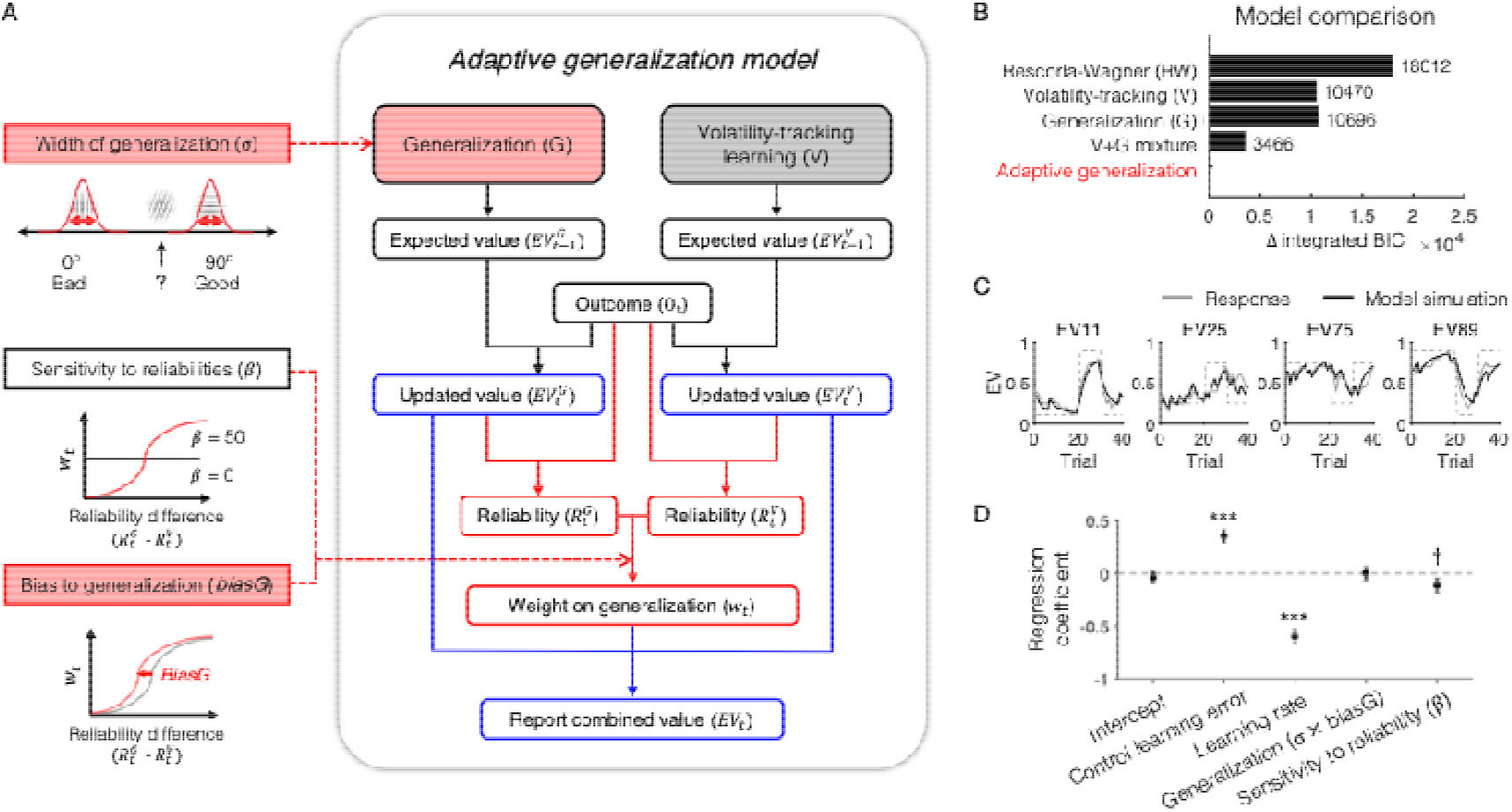
Adaptive generalization model. **(A)** Our purposed model hypothesized that two critical strategies—Volatility-based learning and Similarity-based generalization—operate in tandem. Given that participants were informed of potential volatility in the learning environment, we hypothesized that they would track environmental volatility and incorporate this information to infer the value of stimuli, and that, at the same time, participants would leverage feature similarity to estimate the value of stimulus. The estimated values derived from these two distinct learning strategies are then combined to make choices. The extent to which individuals relied on each strategy was determined by the reliability difference between the two strategies, each reflecting model predictability (i.e., difference between updated value and observed outcome in each trial) estimated based on the corresponding learning strategy. The model introduced two core individual-level parameters that determined the level of generalization; ‘*Width of generalization*’ captured individuals’ tendency to generalize in valuation as a function of feature similarity, and ‘*Bias toward generalization*’ captured individuals’ tendency to favor generalization over the volatility-tracking strategy. A sensitivity parameter ‘*β*’ modulates how strongly the reliability difference influences arbitration. **(B)** Our suggested model best explained participants’ responses during the learning task. This model outperformed alternative models, including Rescorla-Wagner model (RW), Volatility-based value learning model (V), Similarity-based generalization model (G), and mixture of Volatility-based value learning and generalization model (V+G mix). **(C)** Learning curves from an exemplar participant are depicted. Gray lines represent the participant’s actual responses, while black lines depict the simulated responses generated by our suggested model. **(D)** Moreover, participants who exhibited greater sensitivity to reliability differences (*β*) across all four groups tend to show smaller estimation errors even after controlling for baseline learning capacity, learning rate, and generalization tendencies. Error bars represent standard error for standard beta estimates. ^†^*P* = 0.069, \*\*\**P* < 0.001.

This proposed arbitration model (‘V+G arbitration’) explained participants’ estimation responses across all four groups significantly better than a standard Rescorla-Wagner (RW)^23^ reinforcement learning model (t(165) = 26.72, P = 0) and three alternative nested models (**Fig. 2B, 2C, S4A;** see **Methods** for model description). Specifically, formal model comparisons using integrated Bayesian Information Criteria (iBIC) indicated that the V+G arbitration model provided a better fit to participants’ responses than the RW model (ΔiBIC = 18012), the Volatility-tracking model (V; ΔiBIC = 10470), the Generalization model (G; ΔiBIC = 10696), or the model assuming a constant mixture of the two strategies (V+G mix; ΔiBIC = 3466). Consistent with these findings, likelihood-ratio tests also favored the V+G arbitration model over each of the three nested alternatives: V (χ^2^(4) = 10554.70, P < 0.001), G (χ^2^(2) = 10738.39, P < 0.001), and V+G mix (χ^2^(1) = 3487.19, P < 0.001). Moreover, across all four groups, participants who exhibited greater sensitivity to reliability differences (*β*), although the effect was only marginally significant, tended to show smaller estimation errors—measured as the absolute differences between their responses and the true associated reward probabilities, accumulated across all trials (standardized beta = −0.11, SE = 0.060, P = 0.069; **Fig. 2D**; see **Methods**)—even after controlling for baseline learning performance (standardized beta = 0.34, SE = 0.052, P < 0.001), learning rate *α* (standardized beta = −0.60, SE = 0.064, P < 0.001), and generalization tendencies (standardized beta = 0.0023, SE = 0.060, P = 0.97). These results support the hypothesized mechanism that, across different contexts in which prior knowledge varies in usefulness, individuals track both the utility of generalization and changes in environmental uncertainty, and adaptively apply generalization when learning novel associations. Note that all parameters included in the V+G arbitration model including *α, β, σ*, and *biasG* were recoverable and showed high correlations between true and recovered parameter values (*β*: Spearman’s correlation coefficient ρ = 0.55, P < 0.001; *σ*: ρ = 0.86, P < 0.001; *biasG*: ρ = 0.82, P < 0.001; *α*: ρ = 0.90, P < 0.001; see **Methods** for parameter recovery details).

### Individuals are more likely to engage in generalization when the learning environment is generalizable

Among all individuals, we expected those exposed to a generalizable context, i.e., a monotonic stimulus-reward association followed by a positively congruent association for novel stimuli (the mC group), to show the highest reliance on the generalization strategy, provided they employed an adaptive learning approach. Extending our model-agnostic findings from the regression analysis of individuals’ responses during the Stable phase, the two core parameters designed to capture the extent of generalization (*σ* and *biasG*) supported our hypothesis (**Fig. 3A**). Specifically, a two-way ANOVA revealed significant main effects of the type of Anchoring session (m vs. nm) and the type of Generalization session (C vs. IC) on both *σ* (m vs. nm: F(1, 162) = 12.22, P < 0.001; C vs. IC: F(1, 162) = 53.56, P < 0.001) and *biasG* (m vs. nm: F(1, 162) = 6.02, P = 0.017; C vs. IC: F(1, 162) = 15.35, P < 0.001). In addition, the interaction effect was significant for both *σ* (F(1, 162) = 10.68, P = 0.0017) and *biasG* (F(1, 162) = 11.66, P = 0.0011). In contrast to the parameters reflecting individuals’ generalization tendencies, interaction effects were not significant for *α*, the learning rate (C vs. IC: F(1, 162) = 1.23, P = 0.27; interaction: F(1, 162) = 0.67, P = 0.42) or for *β*, the sensitivity to reliability (C vs. IC: F(1, 162) = 0.021, P = 0.88; interaction: F(1, 162) = 1.54, P = 0.22; **Fig. S4B**). The main effects of the type of Anchoring session (m vs. nm) were marginally significant for *α* (m vs. nm: F(1, 162) = 3.90, P = 0.051) and significant for *β* (m vs. nm: F(1, 162) = 4.45, P = 0.037). Specifically, participants who learned monotonic stimulus-reward association tended to have smaller learning rate and greater sensitivity to reliability. These results show that differences in learning context specifically influence the extent to which individuals rely on generalization, and suggest that individuals are more likely to engage in generalization when prior knowledge remains broadly applicable to subsequent learning.

**Figure 3.**
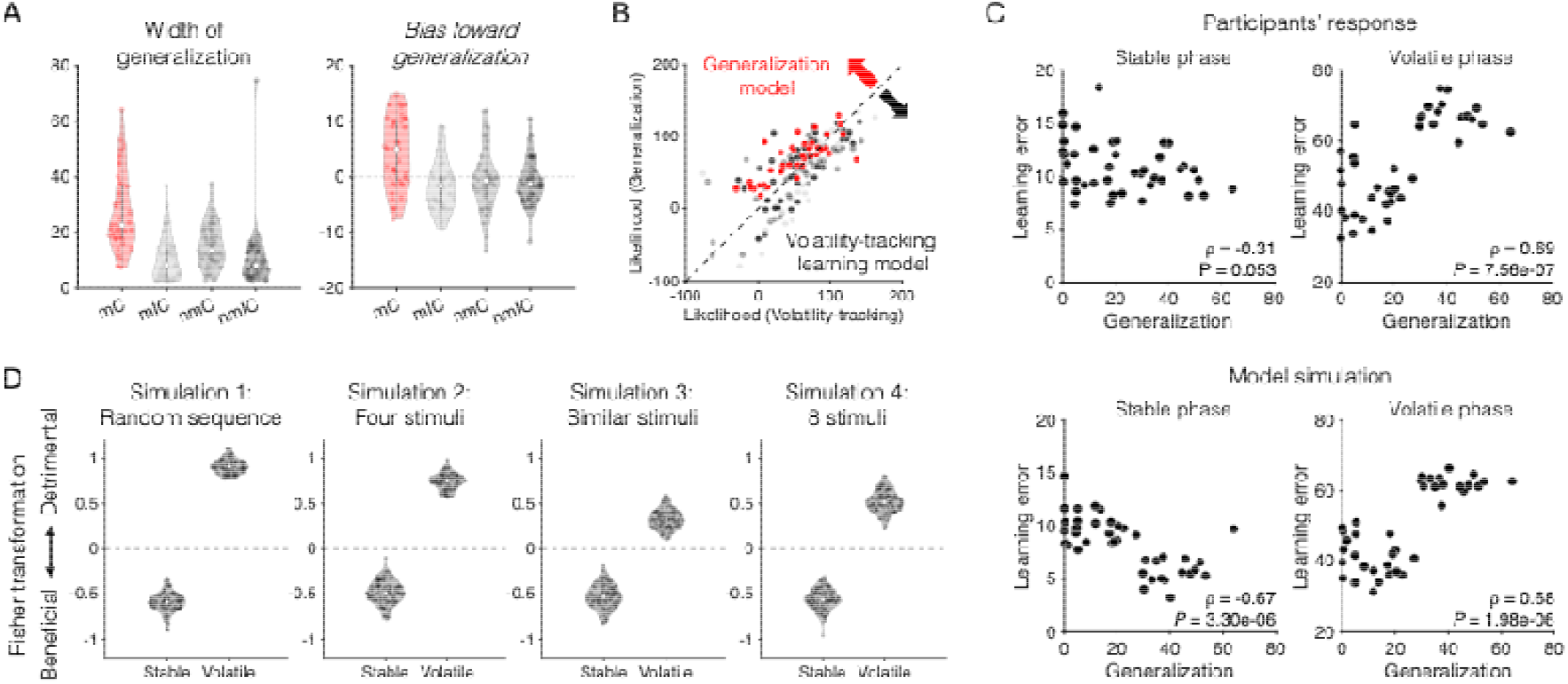
The dominance of generalization strategy in a generalizable context. **(A)** Estimated parameters from our proposed arbitration model. Both generalization-related parameters—width of generalization (σ) and bias toward generalization (*biasG*)—were significantly higher in the monotonic-congruent group (generalizable context) compared to the other three groups (non-generalizable contexts). Learning rates and reliability sensitivity were statistically comparable across all four groups. Error bars represent s.e.m. **(B)** To test whether participants’ responses were biased toward the volatility-based value learning or generalization strategy, we compared model-fits at the individual level. Unlike individuals in other groups (non-generalizable contexts; dark to light gray markers), most participants in the monotonic-congruent group (generalizable context; red markers) exhibited responses that were better explained by the Generalization model rather than the Volatility-tracking value learning model. **(C)** Relying on the generalization strategy facilitated value learning in a stable environment but was detrimental in a volatile environment. **(D)** Our proposed model successfully captured the context-dependent effects of generalization on value learning across the Stable and the Volatile phases. The context-dependent impacts of generalization were replicated across various simulated experimental conditions (simulation n=100). Specifically, (Simulation 1) generalization was beneficial in the Stable phase but detrimental in the Volatile phase when the sequences of presented stimuli and reward outcomes were completely randomized. The context-dependent effects were consistently observed (Simulation 2) when pseudo-subjects were assumed to learn the values of four stimuli simultaneously during the Generalization session, (Simulation 3) when stimuli with similar orientations ([10 and 22.5 degrees] and [67.5 and 80 degrees]) were learned together, (Simulation 4) and when a larger number of stimuli was learned at the same time.

The model-fit differences observed across the four groups between the two single-strategy learning models, each nested within the broader V+G arbitration model, further support the adaptive use of the generalization strategy (**Fig. 3B**). In the mC group, individual-level model-fits favored the Generalization model, which assumes exclusive use of the generalization strategy, over the Volatility-tracking model, which assumes exclusive reliance on the volatility-tracking strategy (t(40) = 4.03, P = 2.44e-04; red markers in **Fig. 3B**). In contrast, responses from participants in the other three groups (mIC, nmC, nmIC) were better explained by the Volatility-tracking learning model or showed comparable model-fits between the two models (mIC: t(43) = −2.07, P = 0.045; nmC: t(40) = −1.36, P = 0.18; nmIC: t(39) = 0.095, P = 0.92; monochrome markers in **Fig. 3B**). These results, in line with the model parameters from our suggested V+G arbitration model, further emphasize that individuals adapt their learning strategies according to the context, depending on the usefulness of prior knowledge.

### Generalization has context-dependent effects on value learning

We further examined how individuals’ biases toward generalization strategy affect individuals’ learning performance. Given that generalization is typically beneficial only when the context is predictable based on prior knowledge, we expected individuals with stronger generalization tendencies to perform better during the Stable phase—when the positive associations between stimulus and orientation is retained—, but worse during the Volatile phase. As expected, there was a negative association between the extent of generalization—measured as the interaction between *σ* and *biasG—* and learning error, defined as the absolute difference between participants’ responses and the true reward probabilities accumulated across all trials in the Stable phase (Spearman’s correlation coefficient ρ = −0.31, P = 0.053), suggesting that generalization was beneficial when prior knowledge remained congruent. However, in the volatile environment, greater reliance on generalization was associated with higher learning errors (ρ = 0.69, P = 7.56e-07), highlighting the detrimental effects of over-generalization on flexible learning (**Fig. 3C**, top). Note that participants’ perceptual sensitivity, as estimated from an independent discrimination task (see **Methods**), was not correlated with their learning errors (ρ = −0.093, P = 0.23) or with their generalization tendencies (*σ* × *biasG*; ρ = 0.095, P = 0.22), suggesting that the extent of generalization cannot be attributed to their perceptual sensitivity.

The context-dependent effects of generalization on value learning were observed not only in the experimental data but also replicated in the data simulated using the V+G arbitration model (Stable phase: Spearman’s correlation coefficient ρ = −0.67, P = 3.30e-06; Volatile phase: ρ = 0.68, P = 1.98e-06; **Fig. 3C**, bottom). One might suspect that the observed differential impacts of generalization were specifically influenced by our experimental design. To test this possibility, we constructed additional hypothetical experimental settings and simulated the behavioral choices participants would have made within each (see **Methods** for simulation details). Specifically, we tested four alternative task designs: 1) a version with fully random stimulus sequences instead of the original pseudo-random sequences; 2) a version in which individuals learned four stimuli within a single session rather than across two separate sessions; 3) a version where two similar Gabor stimuli were learned as a pair within each session; and 4) a version involving the learning of ten stimuli instead of four. In all simulations, stimuli with orientations of 0-, 45-, and 90-degree were used as anchors in the Anchoring session, consistent with the original task paradigm. Across all simulated task settings, the extent of generalization was significantly associated with learning performance. In the Stable phase, this association was significantly negative (Simulation 1: t(99) = −53.07, P < 0.001; Simulation 2: t(99) = −37.32, P < 0.001; Simulation 3: t(99) = −40.05, P < 0.001; Simulation 4: t(99) = −49.84, P < 0.001; **Fig. 3D**), indicating that generalization facilitated learning when task structure was consistent. In contrast, during the Volatile phase, the association was significantly positive (Simulation 1: t(99) = 119.89, P < 0.001; Simulation 2: t(99) = 86.96, P < 0.001; Simulation 3: t(99) = 30.62, P < 0.001;

Simulation 4: t(99) = 45.15, P < 0.001; **Fig. 3D**), suggesting that generalization negatively impacted learning when reward contingencies changed frequently. Both the experimental and simulation results suggest that, regardless of individuals’ ability to adaptively apply learning strategies, excessive reliance on generalization may reduce the efficiency of learning, particularly in unpredictable and rapidly changing environments.

## Discussion

Here, using a probabilistic reward learning task in which stimulus features could be informative, combined with a computational model-based approach, we show that individuals adaptively employ similarity-based generalization during value learning. Specifically, we show that people adjust their reliance on generalization by monitoring its reliability relative to alternative strategies. We suggest that, through this adaptive learning mechanism, individuals tend to generalize more in contexts where prior knowledge benefits subsequent estimation, and less in volatile or unpredictable environments.

Previous studies have shown that information acquired from prior experiences can be generalized to the valuation of novel stimuli^4,24-26^. When stimuli share common feature dimensions, individuals can infer the information about novel stimuli based on feature similarity. In real life, however, environments or contexts change occasionally and unexpectedly, and therefore, previously acquired knowledge does not necessarily remain useful. The current study used a novel probabilistic learning paradigm to directly investigate whether and how individuals adapt to such changes. Our findings extend previous research on individuals’ use of generalization and suggest that people adjust their reliance on generalization based on their beliefs about the usefulness of prior knowledge. Maintained use of a generalization strategy allows individuals to exploit prior knowledge^3,4,27,28^, while reduced use facilitates a trade-off with richer exploration, enabling adaptation to changing environments^29^. This adaptive learning strategy provides a mechanistic explanation for how one can efficiently learn from limited information (e.g., one-shot learning^30,31^).

Historically, research on generalization in humans and rodents has largely focused on the loss domain, particularly in contexts such as avoidance learning and fear conditioning^5-11,13,32-37^, due to the more robust behavioral responses elicited by negative compared to positive outcomes. Such generalization tendencies observed across species can be explained from an evolutionary perspective, as they promote defensive and loss-aversive responses that enhance survival^33,38,39^. A few recent studies, however, have shown that individuals tend to use generalization when making decisions between options associated with positive rewards^4,40-42^, suggesting that the motivation for employing a generalization strategy is not limited to the loss domain but also extends to the gain domain. Given these data, and the particularly pronounced generalization pattern observed in the mC group in the current study, we speculate that—beyond a strong drive to avoid potential losses—generalization may also stem from a broader belief about the statistical properties of the world. To test whether this hypothesis extends to a general mechanism of information processing in the brain, future studies may examine whether individuals’ differing beliefs about the environment (e.g., those shaped by cultural background) predict their tendency to generalize in real-world contexts.

Across a broad range of clinical populations, maladaptive information processing has been repeatedly reported as a core cognitive impairment. For example, individuals with anxiety disorders^6-9,11^, depression^12^, post-traumatic stress disorder (PTSD)^13^, and borderline personality disorder (BPD)^14^ tend to overgeneralize prior knowledge when making decisions, reflecting a failure to calibrate the appropriate level of relational inference between contexts (c.f., individuals with autism spectrum disorder (ASD) undergeneralize^43^). Based on the current findings, we propose that maladaptive behaviors, particularly in learning, may arise from at least two distinct levels of cognitive dysfunction, beyond individuals’ biased beliefs. One involves generalization tendencies that individuals exhibit depending on the context in which they are situated. Although our computational model remained agnostic about the process by which individuals form their initial beliefs about the usefulness of generalization—treating it instead as a trait-like feature—, such context-dependent generalization may still be explained by individuals’ rapid adaptation to changing contexts. The other reflects a failure in the arbitration between learning strategies—a case in which the capacity for generalization may remain intact, but the strategy is used to an inappropriate extent. By adopting our computational perspective on adaptive learning, future studies may help delineate the mechanisms underlying maladaptive behaviors in individuals with psychiatric disorders, potentially involving distinct neurobiological circuits.

Various examples have shown that individuals adapt to given contexts and tend to find ways to make efficient choices^44^. The acquisition and representation of information used in decision-making may reflect contextual distribution^45,46^. When cognitive capacity is limited, individuals tend to adopt simpler and more compressed choice policies^47^. In another case, individuals with unstable or potentially impaired valuation systems relied on alternative heuristic decision strategies^48^ that prioritize clearer and less uncertain information^49^. The current work extends these findings and offers novel insights into the adaptive generalization mechanism via which individuals efficiently learn by exploiting previously acquired information. When previously learned information is deemed reliable, individuals exploit this knowledge by generalizing it to subsequent learning. In contrast, when the information becomes unreliable, they shift back to learning the volatility of the environment. In sum, our findings offer a framework for understanding how different expert systems in the brain interact during value learning. We highlight both the beneficial effects of generalization in predictable environments and the potential adverse effects of overgeneralization in volatile contexts, laying the groundwork for investigating psychiatric disorders in which generalization contributes to maladaptive behavior.

## Methods

### Participants

A total of 251 healthy human participants were recruited. All participants provided written informed consent in accordance with the procedures approved by the Institutional Review Boards of Ulsan National Institute of Science and Technology (UNISTIRB-19-09-A). Of the initial 251 participants, 81 participants were excluded for not meeting the predefined performance threshold during the final three trials of the Anchoring session of the learning task. Specifically, their estimated responses did not fall within the required ranges: 0-10% for the stimulus associated with a 0% reward probability, 90-100% for the 100% reward stimulus, and 10-90% for the 50% reward stimulus. An additional four participants were excluded due to data loss. After applying the exclusion criteria, data from 166 participants (male/female = 107/59, age = 23.25 ± 3.20) were included in subsequent analyses. Participants were randomly assigned to one of four groups at the time of recruitment (see **Learning task**), and the groups used in the final analyses were comparable in the number of included participants (41, 41, 40, and 44), gender distribution (Χ^2^(3) = 0.36, *P* = 0.95), and age (*F*(3, 162) = 0.52, *P* = 0.67; **Table S1**).

### Experimental procedures

The experiment comprised three tasks, completed in a fixed order: a discrimination task followed by two learning tasks. The discrimination task was administered first to ensure that participants could distinguish between Gabor patches with different orientations (**Fig. S2**). In both learning tasks, participants learned reward contingencies associated with seven unique stimuli (**Fig. S1**). In the ‘main’ learning task, the stimuli consisted of Gabor patches with distinct orientations. In contrast, the ‘control’ learning task—designed to assess participants’ baseline learning capacity—used unfamiliar foreign characters from the Amazigh alphabet as stimuli instead of Gabor patches. The order of the main and control learning tasks was counterbalanced across participants. At the end of all tasks, participants received a monetary bonus determined by three factors: the sum of reward outcomes across all trials, performance in the learning tasks—defined as the proportion of trials in which the absolute estimation error (i.e., the difference between the estimated and true probabilities) was less than 10%—, and the number of correctly answered catch trials across both learning tasks. Participants were paid a bonus of up to 17,000 Korean won, in addition to a base compensation of 15,000 Korean won per hour.

### Discrimination task

To evaluate participants’ perceptual ability to discriminate the orientations of Gabor patches, a two-alternative forced-choice discrimination task was administered (**Fig. S2**). Each trial started with a fixation animation in which a circle gradually decreased in size over an average duration of 0.53 seconds (SD = 0.013), designed to orient participants’ attention to the center of the screen. Participants then sequentially viewed two stimuli, each presented for 1500 msec and separated by a 4000 msec interval. After the second stimulus, participants had 2000 msec to respond whether the second stimulus was tilted clockwise or counterclockwise compared to the first, using the right and left arrow keys, respectively. Trials in which participants failed to respond within the time limit were labeled as time-over trials and were appended to the end of the task. Stimulus orientations were selected to span the range used in the main learning task (10°, 22.5°, 67.5°, and 80°). Specifically, 16 Gabor patches were chosen, tilted in 4° increments within two orientation ranges: 4°-32° and 58°-86°. For each range, all possible ordered pairs of distinct orientations, as well as pairs with identical orientations, were included, yielding 64 pairs per range. These 128 pairs were pseudo-randomly shuffled to create stimulus sequences, and one of four pre-generated sequences was randomly assigned to each participant.

Discrimination performance was calculated as the proportion of correct responses across the 128 trials. A softmax function was fitted to participants’ responses to estimate two parameters: an ‘inverse temperature’ (*invTemp*), reflecting participants’ orientation sensitivity, and a perceptual ‘*bias*_*perc*_’ (**Eq.1**):

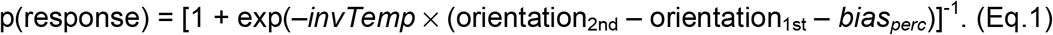

To test whether the four participant groups had comparable perceptual abilities, a two-way ANOVA was conducted on response accuracy and estimated inverse temperature, with the two types of the Anchoring and Generalization sessions (see **Learning task**) entered as main factors. Subsequently, Spearman’s correlation coefficients were computed to examine whether variability in participants’ performance on the main learning task, as well as their generalization tendencies, could be attributed to individual differences in perceptual ability.

### Learning task

We designed a novel probabilistic reward learning task in which participants learned the reward contingencies associated with a set of stimuli, each defined by a unique orientation (‘main’ learning task). The task consisted of an “Anchoring session” followed by a “Generalization session.” A between-group task design was adopted, with participants randomly assigned to one of four groups: 2 types of Anchoring sessions × 2 types of Generalization sessions. In the Anchoring session, participants completed up to 20 trials per stimulus (reward probability = 0%, 50%, 100%). An estimate was considered correct if it deviated by less than 10% from the true reward probability: 0-10%, 40-60%, and 90-100%, respectively. If a participant provided three consecutive correct estimates for each of the three stimuli, they proceeded to the Generalization session, even if they had not completed all 60 trials. In the Generalization session, participants completed 40 trials for each stimulus. The first 20 trials constituted the Stable phase, during which the initial reward contingencies remained unchanged. The next 20 trials made up the Volatile phase, in which the reward contingencies were reversed (i.e., a probability p became *1-p*) and then reversed again after 10 trials. Participants were informed that the reward contingencies might change during the task, although the exact change points were not disclosed.

We designed two types of Anchoring sessions in which the stimulus-reward probability associations were either monotonic or non-monotonic. Specifically, in the monotonic Anchoring session, the associations were set as follows: [0° – 0%], [45° – 50%], and [90° – 100%]. In contrast, the non-monotonic session included [0° – 50%], [45° –0%], and [90° – 100%]. In the subsequent Generalization session, participants learned the reward probabilities associated with four novel stimuli, each defined by a distinct Gabor patch orientation (10°, 22.5°, 67.5°, and 80°). In one of the two types of Generalization sessions, the stimulus-reward associations followed a positive mapping, consistent with the monotonic Anchoring session (‘congruent’ session): [10° – 11%], [22.5° – 25%], [67.5° – 75%], and [80° – 89%]. In the other type, the associations were negative (‘incongruent’ session): [10° – 89%], [22.5° – 75%], [67.5° – 25%], and [80° – 11%]. As noted above, these initial reward contingencies in each type of Generalization session were reversed twice within the session. To reduce cognitive load during the Generalization session, participants were first asked to learn two stimulus-reward associations simultaneously (11 & 89%; or 25 & 75%) and then learn the remaining two. The order of these two parts was counterbalanced across participants within each group to control for order effects. Between the two parts, participants completed the Anchoring session again as a reminder of the assigned stimulus-reward associations (i.e., either monotonic or non-monotonic).

To assess individuals’ baseline learning capacity independent of the stimuli’s feature information, participants completed a ‘control’ learning task in which unfamiliar foreign letters (the Amazigh alphabet) were presented as stimuli instead of Gabor patches. For each participant, the letter stimuli were randomly assigned to specific reward probabilities. The experimental procedure of the control learning task was identical to the main learning task. The order of two learning tasks was counterbalanced across participants in each group to control for order effects.

For both the main and control learning tasks, we generated two types of pre-determined sequences: (1) stimulus presentation sequences and (2) reward feedback sequences. For each Anchoring and Generalization session, four stimulus sequences were generated, ensuring that no stimulus appeared more than twice consecutively. Reward feedback sequences were generated such that, for a stimulus with a reward probability of p%, participants received 10 points on approximately p% of the trials and 0 points on the remaining trials. For stimuli with a 50% reward probability, we ensured that the reward sequence included an equal number of 10-point and 0-point outcomes across the 20 trials in the Anchoring session, with no more two identical reward outcomes appearing consecutively. In the Generalization sessions where two stimuli were learned simultaneously, we generated paired reward sequences, with each pair consisting of one sequence assigned to each stimulus. Reward sequences were independently generated for the two types of Generalization sessions: one congruent and the other incongruent. For each type of Anchoring and Generalization session, one of the four pre-generated stimulus and reward sequences was randomly assigned.

### Linear regression analyses

To examine how prior learning during the Anchoring session influenced subsequent learning in the Generalization session, we conducted regression analyses using participants’ estimates from the Generalization sessions of both the main and control learning tasks (**Fig. 1D**). We first regressed participants’ estimates from both tasks on the true reward probabilities associated with each stimulus. The true reward probabilities had a significant effect on participants’ estimates (β = 0.89, P < 0.001), and the initial model yielded an R-squared value of 0.79, indicating that approximately 79% of the variance was explained. The residuals from this model were then used as dependent variables in a subsequent analysis. Next, to investigate the extent to which participants’ estimates were biased as a function of stimulus feature similarity, we regressed the residuals on a set of independent variables, including group information (2 [monotonic or non-monotonic] × 2 [congruent or incongruent]), stimulus orientations (coded relative to 45°), and their interactions. Specifically, stimulus orientations were coded as −35, −22.5, 22.5, and 35 for the stimuli with 10, 22,5, 67,5, and 80 degrees, respectively, in the main learning task, and as 0 for all stimuli in the control learning task. The regression coefficients associated with this factor (referred to as ‘Anchoring effect’), as well as with the relevant interaction terms, captured the degree of overestimation for the stimuli similar to 90 degrees and underestimation for the stimuli similar to 0 degrees.

To examine the relationship between individuals’ learning performance and estimated model parameters (**Fig. 2D**), we performed robust multiple linear regression using iteratively reweighted least squares with a bisquare weighting function, which reduces the influence of outliers and provides more reliable estimates when the assumption of normality is violated^50^. The dependent variable was the total estimation error in the main learning task. Predictors included the total estimation error in the control learning task, learning rate (α), level of generalization (calculated as the width of generalization multiplied by the bias toward generalization; σ × *biasG*), and reliability sensitivity (β). All variables were z-scored prior to inclusion in the model.

### Computational modeling

To examine the cognitive mechanisms underlying adaptive generalization, we conducted a model-based analysis using participants’ reward probability estimates from the main learning task. Data from the entire learning procedure—including the two Anchoring sessions and the two Generalization sessions—were used to estimate the parameters of the computational models.

#### Rescorla-Wagner model

We used the Rescorla-Wagner model^51^ as the first of a few alternative models to explain participants’ learning behavior. In this model, the reward value of each stimulus is updated according to the reward prediction error, which is calculated as the difference between the observed reward feedback and the predicted value on each trial^23^, with a free parameter—the learning rate α—as follows (**Eq.2**):

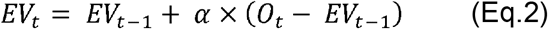

where *t* represents the trial number, *EV* is the estimated reward probability associated with a stimulus, and *O* is the received reward on that trial. This model included one free parameter: the learning rate (α).

#### Volatility-tracking value learning model

We instructed participants that the reward contingencies could change at any time without notice, and thus hypothesized that they would adopt a learning strategy that tracks the volatility of the environment (**Fig. S6**). The volatility-tracking learning model^22^ assumed that participants continuously estimated both the reward probability *r* and the volatility *ν* of the environment, with each evolving according to a Markovian process. Specifically, the probability of reward probability to change from *rt*-1 to rt is assumed to follow a probability distribution centered on the prior expectation *rt*-1, with the width of distribution determined by the volatility *ν* as follows (**Eq.3**):

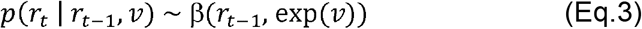

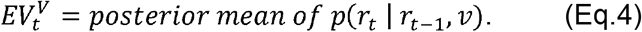

The volatility parameter *ν* modulates how much the observed outcomes can update the estimated reward probability between trials. The reward probability *r* and volatility *ν* is estimated for each unique stimulus and participants are assumed to report the posterior mean of the distribution of the reward probability in each trial (**Eq.4**). Note that the standard beta distribution is parameterized as follows^22^:

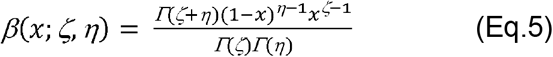

where the expected reward probability = ζ(ζ+η)^-1^, volatility = −log(ζ+η).

Similarly, the model assumed that volatility changed across trials, also as a Markov process as follows (**Eq.6**):

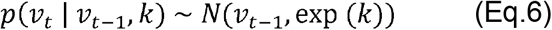

where parameter *k* controls the rate at which volatility changes, defining the width of distribution. A large *k* corresponds to an environment that rapidly switches between stable and volatile states, while a small *k* implies slower shifts in volatility. This hierarchical structure enables the model to adapt its learning dynamically according to changes in the environment.

#### Generalization model

We hypothesized that participants would adopt a generalization strategy based on the shared visual feature of stimulus orientation. Specifically, they might infer the value of previously unencountered stimuli by leveraging their similarity in orientation to those with known reward contingencies. In our main learning task, participants who had learned a monotonic association between stimulus orientations (0°, 45°, and 90°) and reward probabilities (0%, 50%, and 100%) would likely assume that this positive association extends to other novel stimuli with different orientations. To implement this generalization mechanism, we constructed a model that explains the extent to which individuals generalize based on perceptual similarity (e.g., orientation)^4,5^. The core algorithm of this model is that, on each trial, not only the value of the currently presented stimulus, followed by its outcome, is updated based on the prediction error, but also the values of unseen stimuli that share similar feature characteristics (**Eq.7**).

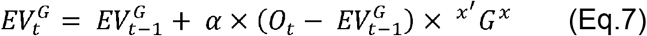

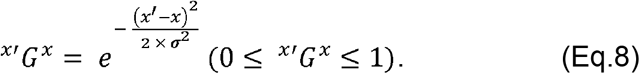

The extent to which each stimulus value is updated is determined by the learning rate (α) and the feature similarity between the presented stimulus (*x*) and all seven stimuli including unseen stimuli (*x*’), modeled as a Gaussian distribution parameterized by the width of generalization (σ) (**Eq.8**). A wider width of generalization indicates broader generalization along the perceptual feature dimension, allowing prediction errors computed for the presented stimulus to influence a broader range of similar stimuli (**Fig. 2A**). For example, a generalization width of 20 yields proportion of generalization (*G*) of approximately 88.25% for an orientation difference of 10°, 53.11% for 22.5°, and 7.96% for 45°. Notably, for the presented stimulus (i.e., *x*’ −*x* = 0), the G term equals 1, indicating a full update of the prediction error. Under a fixed generalization width, proportion of generalization (G) monotonically decreases as feature dissimilarity increases, reflecting diminished generalization to perceptually dissimilar stimuli. This model included two free parameters: the learning rate (α) and the width of generalization (σ).

#### Mixed model between generalization and volatility-tracking

The mixed model assumed that participants’ reward probability estimates (*EV*_*t*_) are predicted by a weighted sum of the estimated reward probabilities from the generalization strategy (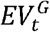; **Eq.7**,**8**) and the volatility-tracking strategy (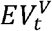; **Eq.3-6**). The two learning strategies are combined using a fixed weight for the generalization strategy (*w*), which ranges from 0 and 1 (**Eq.9**).

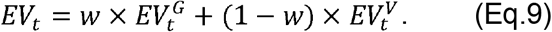

This model included three free parameters: the learning rate (α), the width of generalization (σ), and the weight for the generalization strategy (*w*).

#### Adaptive generalization model

We hypothesized that individuals would adaptively use generalization strategies based on their perceived reliability (**Fig. 2A**). Specifically, the adaptive generalization model assumed that participants evaluate the reliability of each strategy (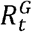 for thegeneralization strategy; 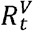for the volatility-tracking strategy), computed based on the estimation error between the observed reward on the current trial (*O*_*t*_) and the expected value up to that trial generated by each learning strategy 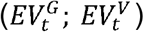

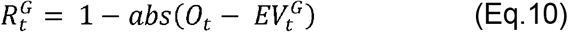

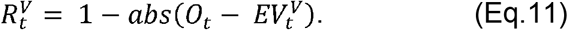

Then, the relative reliability of generalization strategy 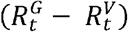 was used to compute the weight (*w*_*t*_) for combining the expected values from the two strategies on trial t (**Eq.12**). Specifically, the weight was determined using a logistic function where the slope *β* reflects sensitivity to the reliability difference and the *biasG* represents the bias toward relying on the generalization strategy:

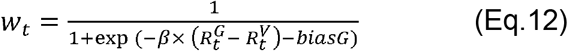

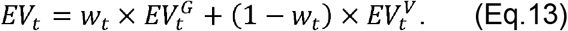

As in the mixed model, the adaptive generalization model assumed that participants’ reward probability estimates (*EVt*) are predicted by a weighted sum of the estimated reward probabilities from the generalization strategy 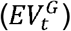 and the volatility-tracking strategy 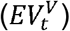 (**Eq.13; Fig. S5-6**). The critical difference in the adaptive model was that the weight was determined on a trial-by-trial basis. This model included four free parameters: the learning rate (α), the width of generalization (σ), the sensitivity to reliability difference (*β*), and the bias toward the generalization strategy (*biasG*). In all statistical analyses, *biasG* was transformed using a sigmoid function, [1 + exp(-biasG)]^-1^, to capture its influence on reward probability estimates.

### Parameter estimation

Each model was constructed using a hierarchical population-level structure, in which individual-level parameters were treated as random effects drawn from group-level distributions. For each parameter, the group-level distributions were modeled as Gaussian, with a standard normal distribution (Normal(0, 1)) and free group-level hyperparameters (mean and standard deviation), following noncentered parameterization^52^. The hyperparameters were estimated using uninformative priors: means ~ Normal(0, 10) and standard deviations ~ Cauchy(0, 2.5) with lower bound of zero. For the learning rate (α) and the sensitivity to reliability difference (β), we applied an inverse probit transformation and then multiplied a constant (1 for α and 50 for β) to restrict the parameters to the ranges [0, 1] and [0, 50], respectively. The bias toward the generalization strategy (*biasG*) was not constrained. The width of generalization (σ) was constrained to be positive by applying an exponential transformation. To estimate the joint posterior distribution of model parameters, we used Markov chain Monte Carlo (MCMC) sampling using the No-U-Turn Sampler (NUTS), a variant of the Hamiltonian Monte Carlo algorithm, implemented in Stan^53^ and accessed through its R interface^52^. For each model, four independent chains were run, each with 5,000 samples, discarding the first 2,000 samples as burn-in. We conducted a visual inspection for convergence and ensured that all values of the potential scale reduction factor (R-hat) were below 1.05 for all variables^54^.

### Model comparison

The adaptive generalization model served as the full model, with all other models defined as its nested variants. To evaluate and compare model fits, we calculated the negative log-likelihood for each model and computed the integrated Bayesian Information Criteria (iBIC) across the entire posterior distribution^55^. In addition, we used likelihood-ratio tests^56^ to examine whether the full model showed better fit over each of the three nested alternatives.

### Parameter recovery

To assess parameter identifiability in the best-fitting model (the adaptive generalization model), we conducted a parameter recovery analysis using simulated data^57,58^. Specifically, we simulated participants’ reward probability estimates using the parameters originally estimated from their empirical data. We then re-estimated the model parameters from the simulated data and computed the Spearman correlation between the original and recovered parameter estimates.

### Estimated parameter comparison

To Investigate how learning contexts influence learning strategies across the four groups (**Fig. 3A, S4B**), we examined the main effects of the Anchoring session type (monotonic vs. nonmonotonic) and the Generalization session type (congruent vs. incongruent). Prior to comparing model parameters across groups, we conducted assumption checks to determine the appropriate statistical test. Specifically, we assessed normality using the Kolmogorov-Smirnov test, and tested for homogeneity of variance using the two-sample F-test. Because at least one of the two assumptions (normality or homogeneity of variance) was violated for all parameters in one or more groups, we used Welch’s two-way ANOVA^59^ for all group comparisons of the estimated parameters from the arbitration model.

### Simulation study

To assess whether the observed context-dependent and adaptive use of generalization across the Stable and the Volatile phases (**Fig. 3D**) was specific to our experimental paradigm, we conducted model-based simulation analyses across four alternative experimental environments. In all simulations, we used the model parameters estimated from participants in the mC group (Anchoring session = monotonic, Generalization session = congruent), allowing us to generate pseudo-behaviors as if the same participants had completed additional experimental conditions. For each alternative setting, we simulated pseudo-behaviors 100 times per agent, using individual parameter set. In all four simulations, agents learned a positive association between stimulus orientation and reward probability during the Anchoring session, followed by the Stable phase of the Generalization session, mirroring the structure of the original experiment conducted with the mC group. Each simulated agent completed 20 trials per stimulus during the Anchoring session.

The first simulation tested the impact of stimulus and outcome sequences on learning. In the original learning task, four pseudo-random sequences for the stimuli and four for the corresponding reward outcomes were pre-generated and randomly assigned to participants. To rule out the possibility that our results were somehow biased from the assigned sequences, the simulation fully randomized both stimulus and outcome sequences across simulated agents, while still adhering to the predefined reward probabilities. Note that, as in the original learning task, the total number of rewarded trials per stimulus was fixed according to its assigned reward probability, but the order of rewarded and unrewarded trials was randomized across simulated agents. The Anchoring session consisted of a fixed number of 20 trials per stimulus for all participants. Aside from the randomization and the fixed number of trials in the Anchoring session, all other task settings remained identical to those in the original task.

The second simulation tested a scenario in which participants learned the reward probabilities of all four stimuli simultaneously within a single Generalization session, rather than across two separate sessions. As in the first simulation, stimulus and outcome sequences were fully randomized across simulated participants, while still adhering to the predefined reward probabilities. All other task settings remained identical to those in the original task.

The third simulation tested the impact of simultaneously learned stimulus pairs. Given that the generalization strategy embedded in the adaptive generalization model updates reward feedback for all stimuli with similar perceptual features, one might expect more pronounced biases when learning stimulus pairs with higher, rather than lower, feature similarity. In this simulation, instead of the original stimulus pairs used in the learning task ([22.5° and 67.5°], [10° and 80°]), a new pair of [10° and 22.5°] was learned within one Generalization session, and a pair of [67.5° and 80°] in another session. The order of the sessions was counterbalanced across simulated participants. As in the first two simulations, stimulus and outcome sequences were fully randomized across simulated participants, while still adhering to the predefined reward probabilities. All other task settings remained identical to those in the original task.

The fourth simulation tested a more cognitively demanding scenario in which participants learned the reward contingencies of eight stimuli (9º, 18º, 27º, 36º, 54º, 63º, 72º, 81º), as opposed to four in the original learning task. These stimuli were positively associated with reward probabilities of 10%, 20%, 30%, 40%, 60%, 70%, 80%, and 90%, and were learned simultaneously during a single Generation session. Stimulus and outcome sequences were again fully randomized as in the other simulations. All other task settings remained identical to those in the original task.

Simulated behaviors were used to compute average learning errors separately for the Stable and Volatile phases within the Generalization session. Pearson’s correlation coefficients were then calculated between the average learning errors and the extent of generalization, defined as the product of the simulated parameters (the width of generalization (σ) and the bias toward generalization (*biasG*)). To assess whether individual correlation coefficients were significantly greater than zero, we first applied Fisher’s r-to-z transformation to approximate a normal distribution. One-tailed one-sample t-tests were then conducted on the transformed values to assess whether their mean was significantly greater or less than zero, depending on the direction of the association observed in the empirical data (a negative association for the Stable phase and a positive association for the Volatile phase).

## Acknowledgements

This work was supported in part by the National Research Foundation of Korea (NRF) (RS-2024-00420674, RS-2025-02263045 to D.C.) and the KBRI basic research program through Korea Brain Research Institute funded by Ministry of Science and ICT (24-BR-03-08 to D.C.).

## Notes

### Competing Interest Statement

The authors have declared no competing interest.

