## Supplementary text for "Adaptive generalization and efficient learning under uncertainty"

Dongil Chung, PhD

(; UNIST, 50 UNIST-gil, Ulsan, South Korea; +82-52-217-2744)

**Table S1. Demographics**

|  | Monotonic-Congruent  (n = 41) | Non-monotonic-Congruent  (n = 41) | Monotonic-Incongruent  (n = 44) | Non-monotonic-Incongruent  (n = 40) | Statistics |
| --- | --- | --- | --- | --- | --- |
| Age [mean ± std] | 23.76 ± 3.36 | 23.02 ± 3.68 | 23.25 ± 2.90 | 22.95 ± 2.85 | *F*(3, 162) = 0.52,  *P* = 0.67 |
| Sex [male/female  (% female)] | 28/13  (31.71%) | 26/15  (36.59%) | 28/16  (36.36%) | 25/15  (37.50%) | χ^2^(3) = 0.36,  *P* = 0.95 |
| Education [n (%)] |  |  |  |  | χ^2^(3) = 12.65,  *P* = 0.18 |
| High school graduate | 2 (4.88%) | 3 (7.32%) | 2 (4.55%) | 1 (2.50%) |  |
| University student | 25 (60.98%) | 26 (63.41%) | 27 (61.36%) | 27 (67.50%) |  |
| University graduate | 1 (2.44%) | 0 (0%) | 7 (15.91%) | 3 (7.50%) |  |
| Graduate enrolled or  completed | 13 (31.71%) | 12 (29.27%) | 8 (18.18%) | 9 (22.50%) |  |

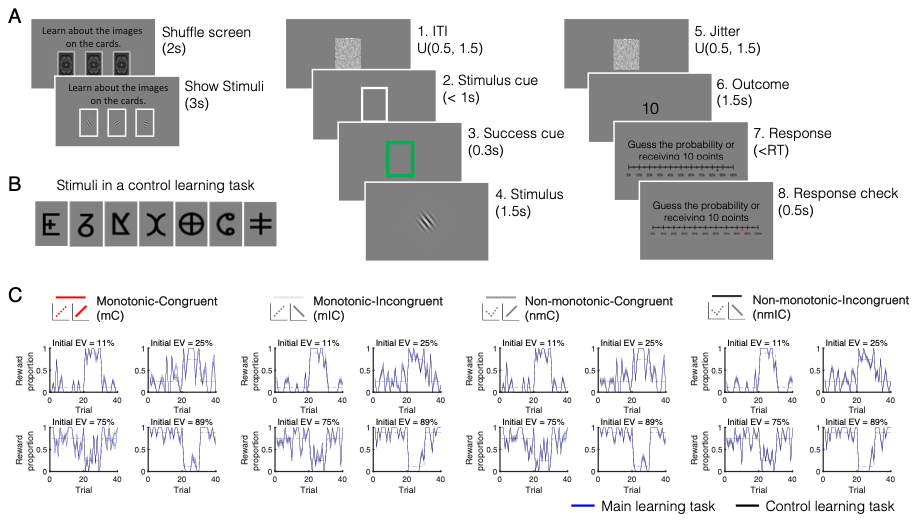

**Figure S1. Experimental paradigm. (A)** Each learning session began with a card-shuffling screen, followed by the presentation of a set of stimuli that would be used during each session. Each trial followed a fixed sequence. At the beginning of the trial, a white noise screen was presented for a randomized duration, followed by a white rectangle prompting participants to press the spacebar within 1 second. A timely response turned the rectangle green, confirming success, and was followed by the stimulus presentation. After a jittered interval, an outcome of either 0 or 10 points was displayed at the center of the screen. Participants then estimated the probability of receiving 10 points for the given stimulus using a 21-point Likert scale ranging from 0% to 100% in 5% increments, adjusted with the ‘<’ and ‘>’ keys and submitted with the spacebar. This trial structure was repeated for a maximum of 20 trials per stimulus in the Anchoring session and 40 trials per stimulus in the Generalization session. During the main and control learning task, participants were occasionally asked to re-enter the reward probability estimate they had reported in the previous trial (not depicted). These catch trials were included up to four times in each Anchoring session and four times in each Generalization session to monitor participants’ attention in both the main and control learning tasks. **(B)** To assess participants’ baseline learning performance independent of feature-based generalization, a control learning task was designed using unfamiliar foreign letters (the Amazigh alphabet), rather than Gabor patches, which were used in the main learning task. **(C)** The control task followed the same trial structure and reward contingencies with the main learning task. Dashed lines represent pre-assigned reward probabilities, black solid lines show average reward proportion per each group in the control learning task, blue solid lines show average reward proportion in the main learning task, and shaded areas indicate s.e.m.

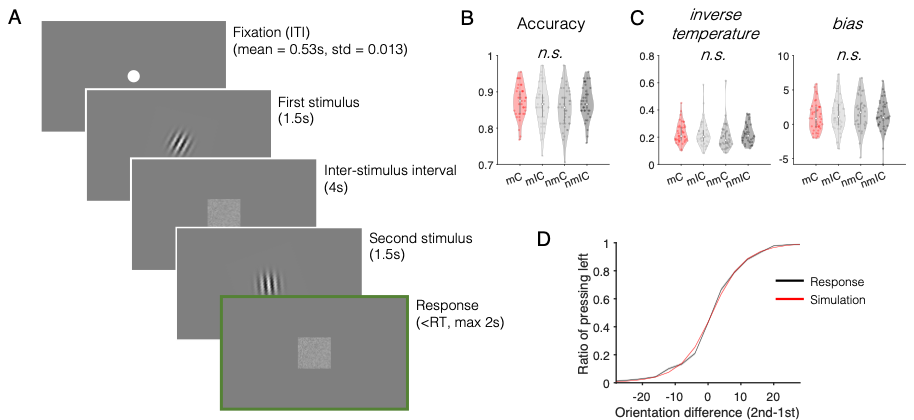

**Figure S2. Discrimination task. (A)** Each trial began with a fixation animation in which a circle gradually decreased in size, guiding participants’ attention to the center of the screen. Participants then viewed two Gabor patch stimuli sequentially, separated by a 4-second inter-stimulus interval. Following the second stimulus, participants responded whether the second stimulus was tilted clockwise or counterclockwise relative to the first one. **(B)** Accuracy was calculated as the proportion of correct responses across 128 trials. All four groups showed above chance performance (0.5). Performances among the four groups were comparable, such that both the main effects of Anchoring session type (monotonic (m) vs. non-monotonic (nm)) and Generalization session type (congruent (C) vs. incongruent (IC)), as well as the interaction effect were statistically not significant (m vs. nm: F(1, 162) = 0.76, P = 0.38; C vs. IC: F(1, 162) = 0.36, P = 0.55; interaction effect: F(1, 162) = 3.19, P = 0.076). **(C)** Perceptual sensitivity (inverse temperature) and bias parameters were estimated using a softmax model fitted to participants’ response data. Neither the main effects (Anchoring session type and Generalization session type) nor the interaction effect was significant for the perceptual sensitivity parameter (m vs. nm: F(1, 162) = 1.15, P = 0.29; C vs. IC: F(1, 162) = 0.39, P = 0.53; interaction: F(1, 162) = 1.27, P = 0.26), as well as for the bias parameter (m vs. nm: F(1, 162) = 0.44, P = 0.51; C vs. IC: F(1, 162) = 0.0084, P = 0.93; interaction: F(1, 162) = 1.26, P = 0.26). **(D)** Model-based simulated data were well matched to participants’ responses on average. Black line represents average participants’ response, red line represents average simulated response based on the computational model, and shaded areas indicate s.e.m.

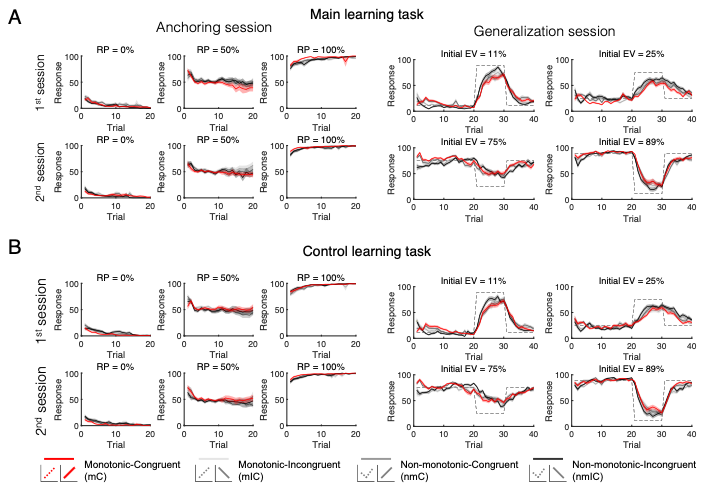

**Figure S3. Learning curves in the main and control learning tasks. (A,B)** In both the main learning task, where Gabor patches were used as stimuli, and the control learning task, which used unfamiliar foreign letters, participants successfully learned the assigned reward probabilities for each stimulus**.** Dashed lines represent the true reward probabilities, solid lines indicate the average estimated probabilities across participants, and shaded areas indicate the s.e.m. Note that the number of trials in the Anchoring session varied across participants depending on their performance, resulting in varying sample sizes across trials when computing the average responses.

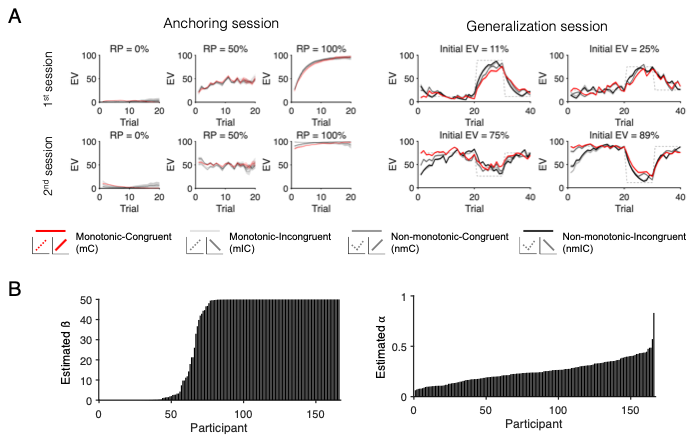

**Figure S4. Simulated learning curves in the main learning task and estimated individual-level model parameters from the adaptive generalization model. (A)** The learning curves depict simulated responses for each of the seven unique stimuli, generated using the adaptive generalization model with parameters estimated from participants’ behavior in the main learning task. Dashed lines represent the true reward probabilities, solid lines indicate the average estimated probabilities across participants, and shaded areas indicate the s.e.m. Note that the number of trials in the Anchoring session varied across participants depending on their performance, resulting in varying sample sizes across trials when computing the average responses. **(B)** Among the estimated individual-level parameters from the adaptive generalization model, β, the sensitivity to reliability, and α, the learning rate, are displayed in ascending order.

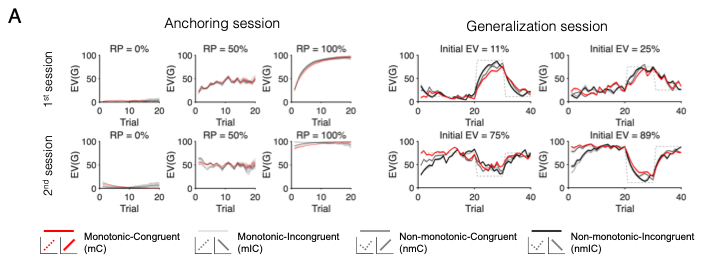

**Figure S5. Simulated learning curves based on the generalization strategy in the main learning task.** The adaptive generalization model includes two learning strategies and their arbitrated use. To illustrate the contribution of the generalization strategy in the learning performance, we generated simulated responses based on the generalization strategy using the parameters estimated from the winning model (i.e., adaptive generalization model) fitted to participants’ behavior in the main learning task. The mC group (Anchoring = monotonic, Generalization = congruent; red lines) stood out by exhibiting superior performance during the Stable phase and inferior performance during the Volatile phase, a pattern that appears to result from their greater reliance on the generalization strategy compared to the other groups. Dashed lines represent the true reward probabilities, solid lines indicate the average estimated probabilities across participants, and shaded areas indicate the s.e.m. Note that the number of trials in the Anchoring session varied across participants depending on their performance, resulting in varying sample sizes across trials when computing the average responses.

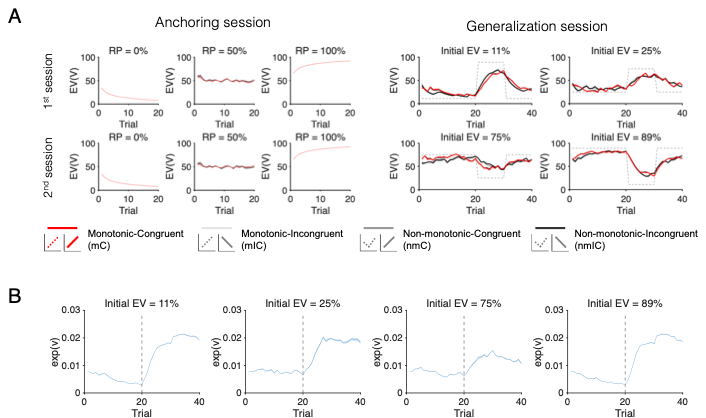

**Figure S6. Simulated learning curves based on the volatility-tracking learning strategy in the main learning task. (A)** To illustrate the contribution of the volatility-tracking learning strategy in the learning performance, we generated simulated responses based on this strategy. Dashed lines represent the true reward probabilities, solid lines indicate the average estimated probabilities across participants, and shaded areas indicate the s.e.m. **(B)** The volatility for each stimulus during the Generalization session was depicted on a trial-by-trial basis. Volatility estimates decreased over time during the Stable phase but increased during the Volatile phase. Solid lines indicate the average estimated volatility across participants, and shaded areas indicate the s.e.m.
